# A generalised approach to detect selected haplotype blocks in Evolve and Resequence experiments

**DOI:** 10.1101/691659

**Authors:** Kathrin A. Otte, Christian Schlötterer

## Abstract

Shifting from the analysis of single nucleotide polymorphisms to the reconstruction of selected haplotypes greatly facilitates the interpretation of Evolve and Resequence (E&R) experiments. Merging highly correlated hitchhiker SNPs into haplotype blocks reduces thousands of candidates to few selected regions. Current methods of haplotype reconstruction from Pool-Seq data need a variety of data-specific parameters that are typically defined ad hoc and require haplotype sequences for validation. Here, we introduce haplovalidate, a tool which detects selected haplotypes in a broad range of Pool-seq time series data without the need of sequenced haplotypes. Haplovalidate makes data-driven choices of two key parameters for the clustering procedure, the minimum correlation between SNPs constituting a cluster and the window size. Applying haplovalidate to simulated and experimental E&R data reliably detects selected haplotype blocks with low false discovery rates – independent if few or many selection targets are included. Our analyses identified an important restriction of the haplotype block-based approach to describe the genomic architecture of adaptation. We detected a substantial fraction of haplotypes containing multiple selection targets. These blocks were considered as one region of selection and therefore led to under-estimation of the number of selection targets. We demonstrate that the separate analysis of earlier time points can significantly increase the separation of selection targets into individual haplotype blocks. We conclude that the analysis of selected haplotype blocks has a large potential for the characterisation of the adaptive architecture with E&R experiments.

## Introduction

Experimental evolution combined with whole genome re-sequencing (E&R) is an effective approach to detect genomic signatures of adaptation (Turner *et al.*, 2013). E&R studies on complex sexually reproducing organisms like *Drosophila* use polymorphic founder populations and selection acts mainly on standing genetic variation instead of new mutations (Tenaillon *et al.*, 2012). Here, the power to detect selected SNPs significantly increases with an increasing number of replicates (Kofler and Schlötterer, 2014; Long *et al.*, 2015) and time points (Burke *et al.*, 2014), and the most economic approach is to sequence pools of individuals (Pool-seq) instead of individual genomes (Schlötterer *et al.*, 2014). While estimating population allele frequencies accurately, Pool-seq does not provide linkage information. Therefore, E&R studies typically focus on individual SNPs instead of incorporating the underlying haplotype structure, and frequently report an excess of outlier SNPs responding to selection (Burke *et al.*, 2010; Turner *et al.*, 2011, 2013; Orozco-terWengel *et al.*, 2012; Remolina *et al.*, 2012; Tobler *et al.*, 2014; Jha *et al.*, 2015; Griffin *et al.*, 2017; Barghi *et al.*, 2019), which is not compatible with population genetic theory (Nuzhdin and Turner, 2013).

Franssen *et al.* (2015) shed some light on the excess of candidate loci by jointly analysing Pool-seq data and experimentally phased haplotypes from the same experiment. They pointed out that a high number of the candidate SNPs in *Drosophila melanogaster* studies were either located in large segregating inversions (Kapun *et al.*, 2014) which suppress recombination or in genomic regions with reduced recombination rates. Another factor contributing to the large number of candidate SNPs is selection on low frequency alleles. The moderate number of recombination events during the experiment is not sufficient to break up the association between the target of selection and linked neutral variants that were private to the selected low-frequency haplotype. These results show that understanding the genomic architecture of adaptation is a very challenging task and individual haplotypes from evolved populations greatly facilitate it by providing linkage information.

Apart from experimental phasing, which requires living flies (Langley *et al.*, 2011; Franssen *et al.*, 2015), various methods have been suggested to statistically infer haplotypes from Poolseq data. Taking advantage of sequenced founder haplotypes, the haplotypes of evolved individuals have been determined by regression (Long *et al.*, 2011), a hidden Markov-model (Cubillos *et al.*, 2013), maximum likelihood (Kessner *et al.*, 2013), and a system of linear equations (Cao and Sun, 2015). These methods rely on the complete knowledge of all involved founder haplotypes (Cubillos *et al.*, 2013) and are limited to a restricted window size because otherwise the error-rate is too high (Long *et al.*, 2011; Kessner *et al.*, 2013; Cao and Sun, 2015).

A different approach to reconstruct selected haplotype blocks without information about the founder haplotypes was proposed by Franssen *et al.* (2017). This approach uses window-based correlation analysis of allele frequency data across replicates and time points combined with hierarchical clustering. Each cluster of SNPs corresponds to a selected haplotype block. Franssen *et al.* (2017) focused on haplotype blocks starting from low allele frequencies (*≤* 0.03), and marker SNPs which are mostly private to them can be identified by strongly correlated allele frequency changes. This approach successfully identified selected haplotype blocks up to several Mb in simulated and empirical Pool-seq data. Extending the approach of Franssen *et al.* (2017) to haplotypes by including also alleles with higher starting frequencies, Barghi *et al.* (2019) successfully reduced over 50,000 outlier SNPs to 99 reconstructed haplotype blocks responding to selection in experimentally evolved *Drosophila simulans* populations. Both, Franssen *et al.* (2017) and Barghi *et al.* (2019) relied on experimentally phased haplotypes of evolved populations to validate their results. Without sequences of evolved and ancestral haplotypes, the validation of reconstructed blocks is challenging, as haplotype reconstruction requires ad hoc choices of key parameters which can change the outcome dramatically and are highly dependent on the data set.

Here, we propose a new approach to define the haplotype reconstruction criteria, which avoids ad hoc choices of clustering parameters and does not depend on the availability of phased haplotype data. Our approach takes advantage of the full genomic data to distinguish between statistically significant clustering, most likely caused by directional selection, and random associations. It is implemented in the R package haplovalidate.

## New Approaches

Haplovalidate is an extension of the R package haploReconstruct (Franssen *et al.*, 2017), which uses hierarchical clustering on pairwise correlations of SNP allele frequencies at multiple time points. A cluster of correlated SNPs in a genomic window defines a reconstructed haplotype block that increases in frequency over time. HaploReconstruct requires a variety of parameters for haplotype detection, which are typically defined ad hoc and tailored to detect selected haplotypes starting at low frequency. Haplovalidate overcomes these limitations by estimating key clustering parameters from the full genomic data.

### Conceptual Idea

The cut-off for determining whether SNPs are clustering or not clustering has a major influence on haplotype reconstruction; too stringent clustering splits regions that belong to the same haplotype block and too relaxed clustering combines independent regions into haplotype blocks (Franssen *et al.*, 2017). Haplovalidate addresses this by generating haplotype blocks for different SNP correlation cut-offs. The approach aims to identify the correlation cut-off which combines blocks that are not independent, but also separates independent blocks. The challenge is to determine which blocks are independent. This is achieved by comparing the correlation of SNPs in the focal cluster with SNPs of other clusters on the chromosome (focal cluster correlations) and SNPs in the focal cluster with SNPs from clusters on the other chromosome (background cluster correlation). With no physical linkage between different chromosomes, background cluster correlations are an estimate of random associations.

Each set of haplotype blocks (with a given correlation cut-off) is tested for the independence of its clusters. Allele frequency trajectories generated by the same selection target should be highly correlated even when separated into different clusters by a too stringent SNP correlation cutoff. If this is the case, focal cluster correlations are significantly higher than the background cluster correlations. If there is no significant difference between focal and background correlations, we assume that all clusters on the focal chromosome are independent and represent different selected regions.

### Haplotype Reconstruction Parameters

For the reconstruction of haplotypes using haploReconstruct (Franssen *et al.*, 2017) at least eight input parameters are needed. Haplovalidate estimates the key parameters from the data, while others are not modified, but were chosen to fit to a broad set of data (see table 1).

**Table 1:** haploReconstruct parameters and the values used for haplovalidate.

| Reconstruction Parameter | Definition | Haplovalidate Value |
| --- | --- | --- |
| **Variable parameters** |  |  |
| min.cl.cor | minimum correlation threshold for SNP clustering | set by haplovalidate |
| **Fixed parameters** |  |  |
| winsize | window size | MNCS of 0.01 or 0.03 |
| min.minor.freq | minimum starting allele frequency of SNPs used for clustering | 0 |
| max.minor.freq | maximum starting allele frequency of SNPs used for clustering | 1 |
| minfreqchange | minimum frequency change of SNPs used for clustering | 0 |
| minrepl | minimum number of replicates in which SNP is present | 1 |
| min.cl.size | minimum number of SNPs in a cluster | 20 |
| min.inter | minimum number of SNPs that overlap for merging clusters | 4 |

### Fixed parameters

We fixed the haplotype reconstruction parameters such that alleles starting from any frequency (starting allele frequency between 0 and 1) in at least 1 replicate are included. The allele frequency change threshold parameter aims to focus on SNPs changing more than expected under drift. Because the *χ*^2^-test and CMH-test used here already account for drift (Spitzer *et al.*, 2019) we set the haploReconstruct threshold to 0. Following Barghi *et al.* (2019), we required at least 20 SNPs for each cluster and only clusters sharing at least 4 SNPs could be merged.

### Variable parameter

In addition to the SNP correlation cut-off, we also varied the window size to fine-tune haplovalidate, as window-size determines the maximal length of reconstructed haplotype blocks. Small windows result in shorter blocks (Franssen *et al.*, 2017), which facilitates independent block separation. Hence, while it is preferred to have small window sizes, this may result in too few SNPs in a window for a reliable estimate of the cluster correlations.

As the landscape of a Manhattan plot can differ dramatically between experiments, the same window-size is not suitable for all data. To use a window-size accounting for the peak landscape we calculate the *median normalised cmh-score sum (MNCS)* as the median cmh-score sum per window normalised by the sum of all cmh-scores (see equation 1). We use this parameter to automatically choose the window-size. Window sizes were varied from 0.1 to 10 Mb in 0.1 Mb steps and the MNCS was calculated for each window-size.

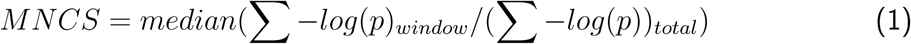

Standard haplovalidate analyses use a window-size corresponding to a MNCS of 0.01. In the case that data-sets do not produce a sufficient number of clusters because the windows were too small we used a MNCS of 0.03 and the corresponding window-sizes.

### Detailed procedure

Haplovalidate consists of four different steps (see figure 1).

**Figure 1:**
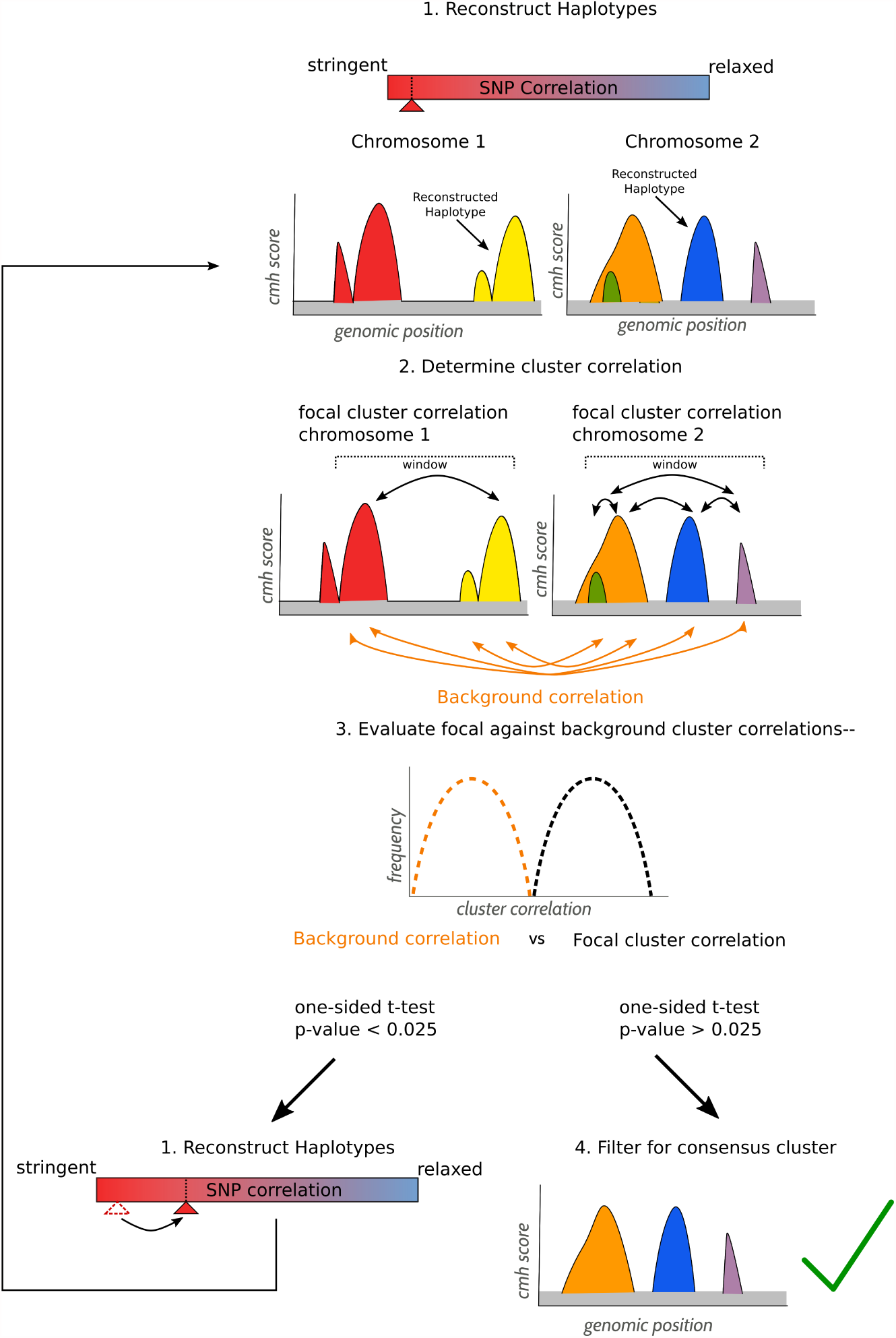
Overview of the iterative procedure to define haplotype blocks with haplovalidate. The haplotype reconstruction starts with stringent parameters (step 1). The correlation of SNPs from different clusters on the same chromosome (focal cluster correlation) and are compared to the correlation of SNPs from clusters located on different chromosomes tested for significant differences (step 2, 3). If focal cluster correlations are higher than the background correlations, this indicates that a too stringent correlation cut-off was used and haplotype reconstruction is repeated using less stringent parameter (back to step 1). If focal cluster correlations are similar to background correlations, the last significant haplotype reconstruction is used and regions with overlapping clusters are filtered for the most dominant cluster (step 4).

**Step 1**: Haplotypes are reconstructed with minimum cluster correlations ranging from 0.9 to 0.3 in 0.1 steps. The reconstruction with most clusters is used as starting point to determine focal and background cluster correlations.

**Step 2**: We normalise allele frequency data by using arcsine-square-root transformation followed by centring and scaling. Cluster correlations are computed by calculating the distribution of median allele frequency trajectory correlations between two clusters. These two clusters can be either clusters within the window used for haplotype reconstruction (focal cluster correlations) or clusters on different chromosomes (background cluster correlations). Focal cluster correlations can be only calculated if at least two clusters are present in a given window. Windows containing only one cluster are not considered. For clusters with more than 2000 SNPs only 2000 randomly selected SNPs were used to increase computational efficiency.

**Step 3**: Cluster correlations are normalised using the Fisher transformation (Fisher, 1915). The difference in focal and background correlations is determined by a one-sided t-test. In the case of a significant difference between chromosome and background (p-value *<* 0.025), the procedure is repeated with step 1 and a less stringent minimum cluster correlation (0.01 steps). If there is no significant difference between chromosome and background (p-value *≥* 0.025), haplovalidate uses the last significant haplotype reconstruction, therefore reducing non-independent haplotype blocks to a minimum. For significance testing cluster correlations and background correlations are used for all windows. If fewer than 3 values are available, haplovalidate returns no result.

**Step 4**: If a selection target is present in several haplotypes, they will be identified as independent, overlapping clusters. To identify the dominating cluster per selection target, we filter genomic regions with overlapping cluster for the cluster with the most significant allele frequency change (cmh-test/*χ*^2^-test, Spitzer *et al.* (2019)).

### Multiple-target haplotypes

Selection targets that share a haplotype have highly correlated allele frequency trajectories (e.g. orange and pink line in figure 2) as the haplotype block behaves as a (additive) single target of selection. Depending on the number and frequency of the selected SNPs in the founder population, it is possible that a given selection target can be found on haplotypes with a variable number of additional selected SNPs. Haplotypes with multiple selection targets are also created by recombination during the experiment. With additive fitness effects, haplotypes with multiple selection targets are favoured and out-compete haplotypes with fewer selection targets (compare top and bottom panel in figure 3). This creates complex allele frequency trajectories which are dominated by the multiple target haplotypes at later time-points. For earlier time-points, when multiple target haplotypes are not yet dominant, the allele frequency changes of single selection targets haplotypes can still be distinguished. The comparison of earlier and later time-points provides therefor the opportunity to disentangle multiple targets on a haplotype (see figure 3 top right plot).

**Figure 2:**
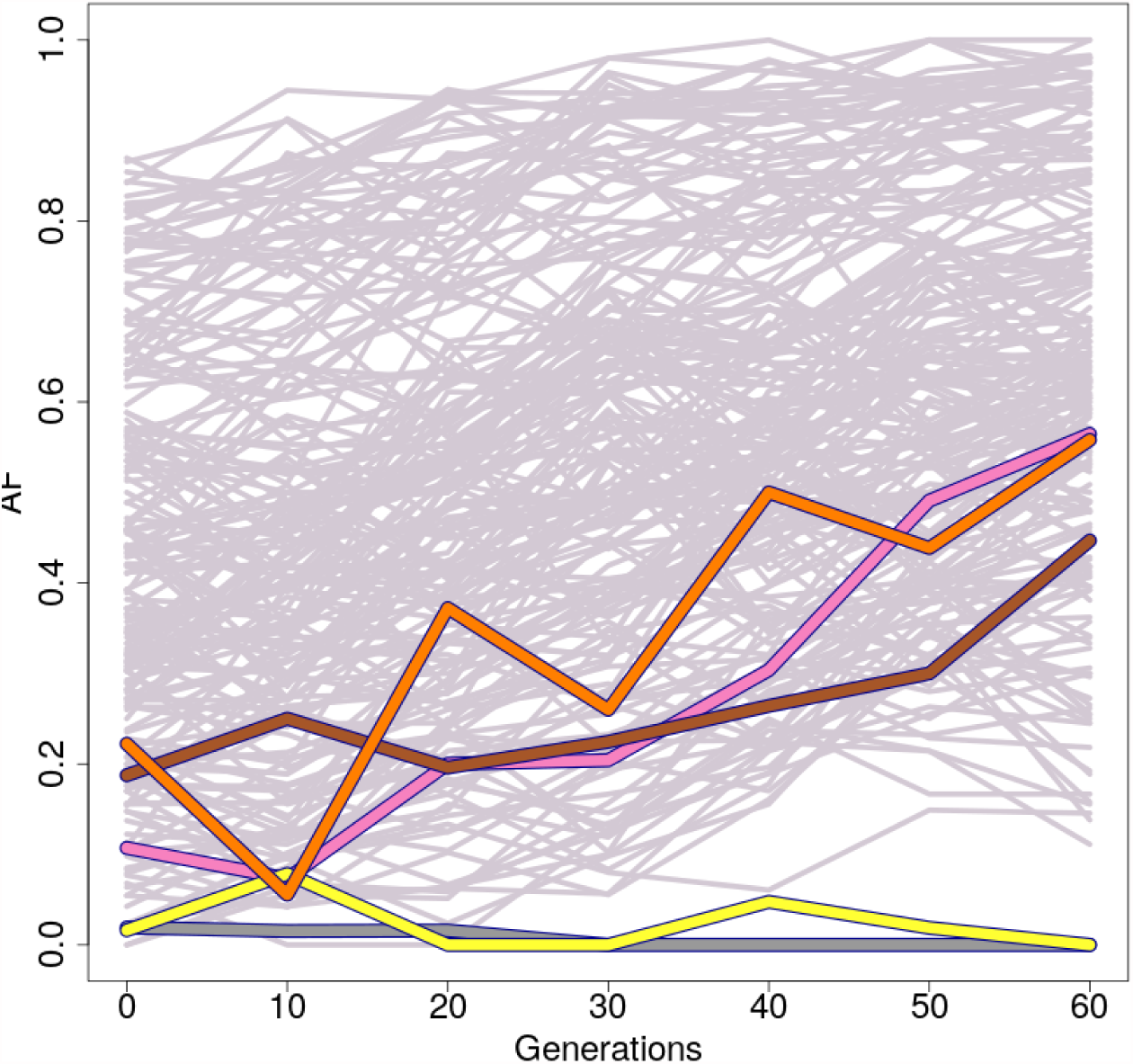
Allele frequency trajectories of a reconstructed haplotype block in one replicate over 60 generations. Selection targets in the region are represented by coloured lines.

**Figure 3:**
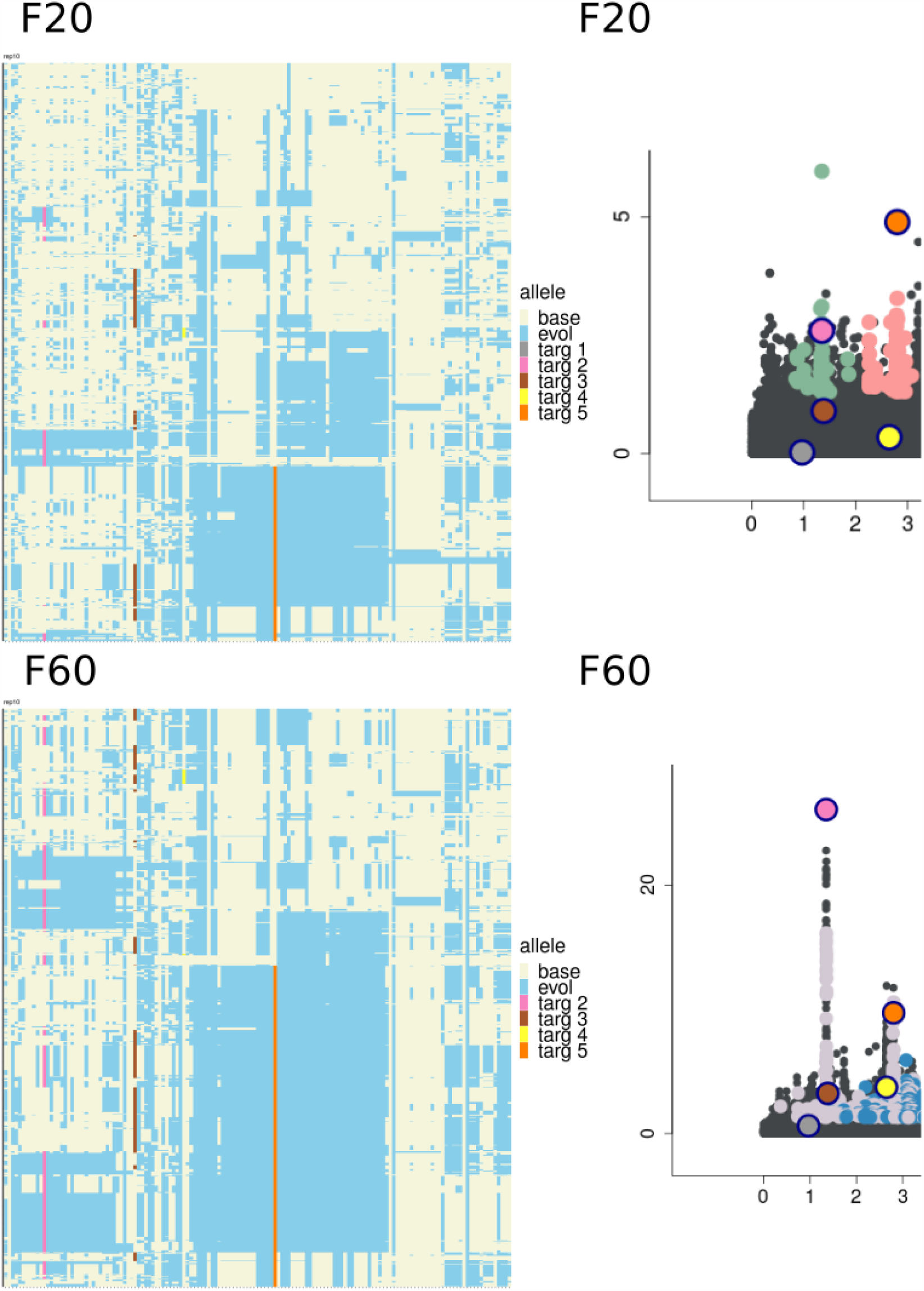
Haplotype blocks containing several selected SNPs behave as single target of selection in the same region as figure 2). Left: Haplotypes present in F20 (top) and F60 in one replicate of a region containing five targets of selection. Each line represents a haplotype and each column indicates the genomic position. Alleles increasing in frequency are marked in grey (colours for the selected SNPs), decreasing alleles are white. Right: Manhattan plot of the haplotype blocks detected by haplovalidate in generation 20 (top) and generation 60 (bottom). Whereas the analysis of F60 results in one region, generation of F20 results in two separate regions, which correspond to haplotype groups present at these time points. Selected SNPs show the same colours as in plot 2 and in the left panel.

## Results

### Haplovalidate Performance

We benchmarked haplovalidate using 1000 selective sweep simulations on two entire chromosomes of *D. simulans* with different numbers of selection targets (16, 32, 64 and 128; 250 simulations each). 13 simulations containing 16 targets, 3 simulations containing 64 targets and 31 simulations containing 128 targets did not result in a sufficient number of clusters to run haplovalidate, therefore we excluded them from the analysis. Haplovalidate reliably detects haplotype blocks containing selection targets as 93 – 98 % of all target SNPs were captured (median values, see figure 4 A). However, not every target SNP resulted in an independent cluster – only 7 % to 56 % of the target SNPs did so (figure 4 B) because many inferred haplotype blocks contain more than one target of selection (see figure 3). The fraction of haplotype blocks that have several targets of selection can be substantial (median 38 % – 91 %, see figure 4 D) with 7% – 14 % of the targets located on a single haplotype block (see figure 4 E). The false positive rate was low, ranging from 4 to 14 % (figure 4 F).

**Figure 4:**
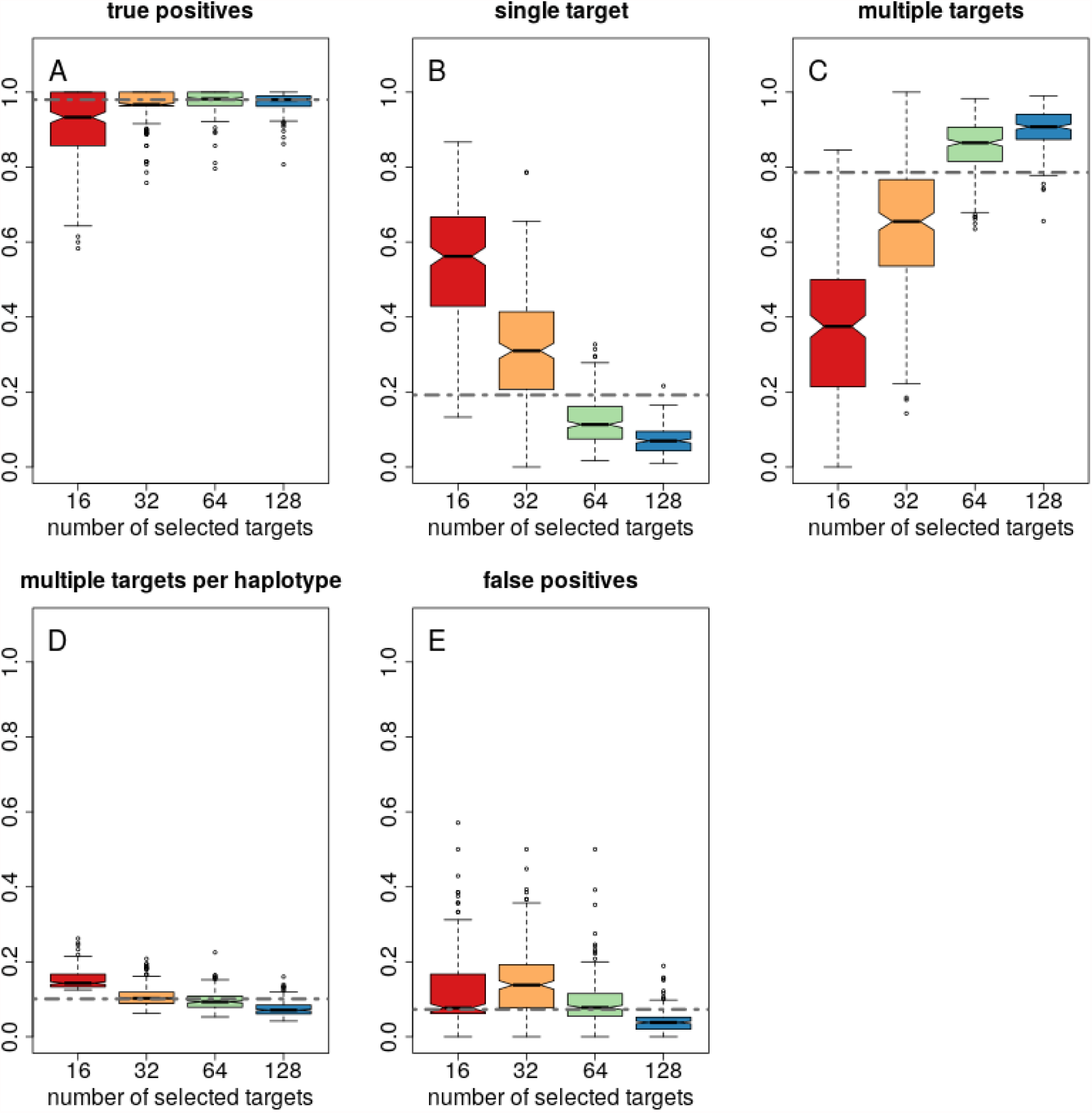
Haplovalidate performance evaluated on 1000 selective sweep simulations containing either 16, 32, 64 or 128 selected alleles. A: True positive rate, B fraction of selected haplotype blocks with single targets, C fraction of selected haplotype blocks with multiple targets, D average number of selection targets per haplotype block, E false positive rate. Dashed line represents the overall median for each parameter.

Haplotype blocks containing multiple targets of selection can either be present at the beginning of the experiment or emerge in later generations due to a recombination event. Using the founder haplotypes, we can determine whether haplotype blocks containing multiple targets were already present in the founder population.

Across all simulations 37,846 targets SNPs (82 %) were located on a haplotype with other targets SNPs at generation 60. 95 % of these targets share haplotypes already in the founder population. This is significantly more often than random pairs of 35,000 SNPs having the same distances (*χ*^2^-test, p-value *<* 0.001). This result indicates that haplotype structure in the founder population pre-determines the occurrence of shared haplotype blocks.

### Fine mapping of multiple-target haplotype using intermediate time-points

Intermediate time-points can be only informative when a sufficient number of candidate SNPs is available for clustering. 74 simulations with 32 targets, 166 with 64 targets and 126 with 128 targets contained sufficient candidate SNPs for the analysis of intermediate time-points.

The inclusion of intermediate time-points increased the number of haplotype blocks while decreasing the average number of selection targets per haplotype block (see figure 5). This was also observed when simulations with different numbers of selection targets were analysed separately (see supplementary data S1).

**Figure 5:**
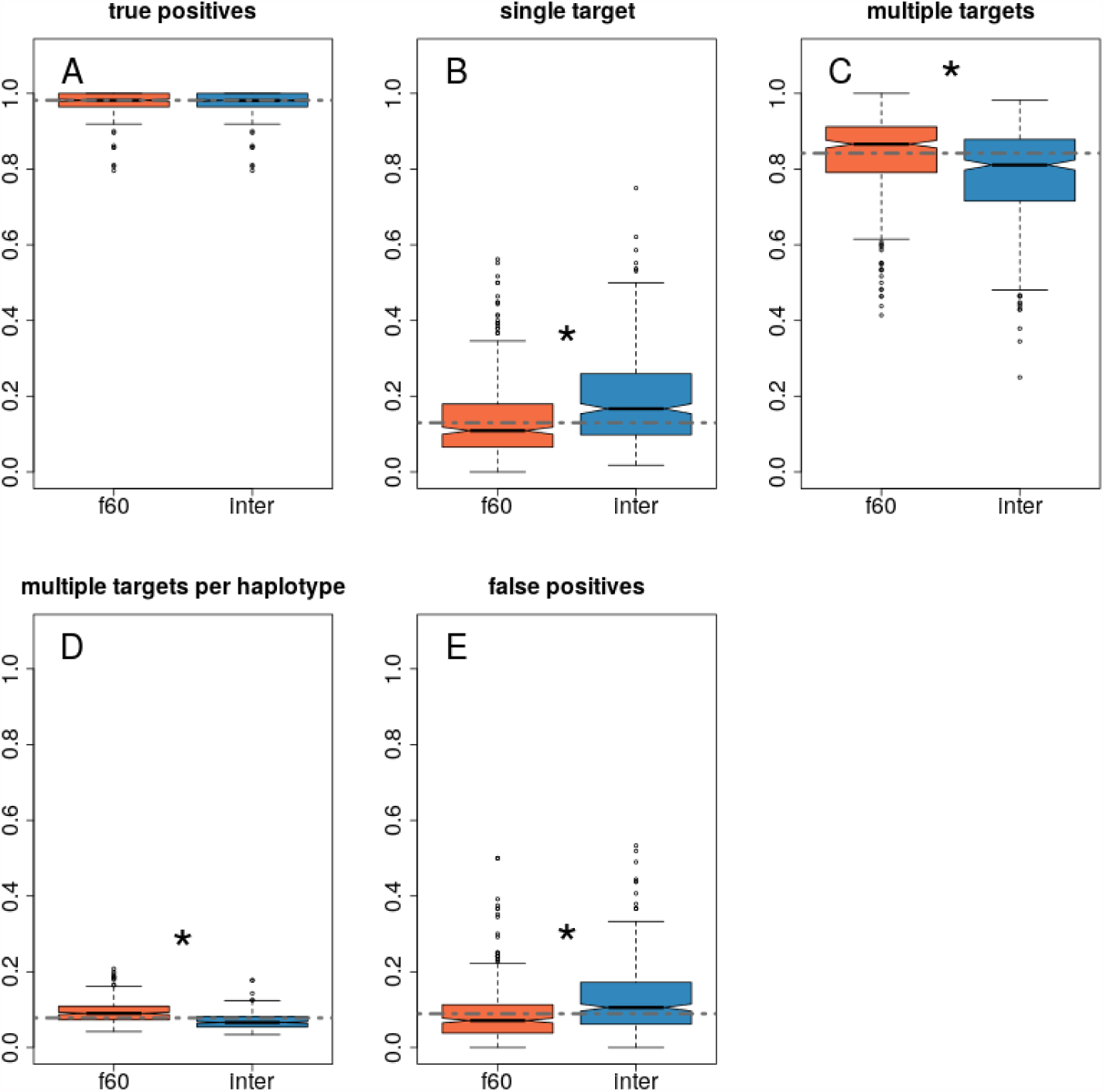
Haplovalidate with and without including intermediate time-points. A: True positive rate, B: fraction of selected regions with a single selection target, C: fraction of haplotype blocks with multiple targets, D average number of selection targets on a haplotype block, E: false positive rate. Dashed line represents the overall median for each parameter. Asterisks indicate significant differences for clustering with and without intermediate time-points (p-value < 0.05).

Using only intermediate time-points resulted in 32 % of the clusters not containing a selection target (see supplementary data S2). 74 % of these clusters are based on hitchhiking SNPs connected to a target of selection, but while the hitchhikers passed the significance threshold, the causative SNP did not and was therefore not included in the clustering. Hence, even when the causative SNP is missing from the candidate SNPs, the correct selected region can be already identified in early generations (for an example see figure 6).

**Figure 6:**
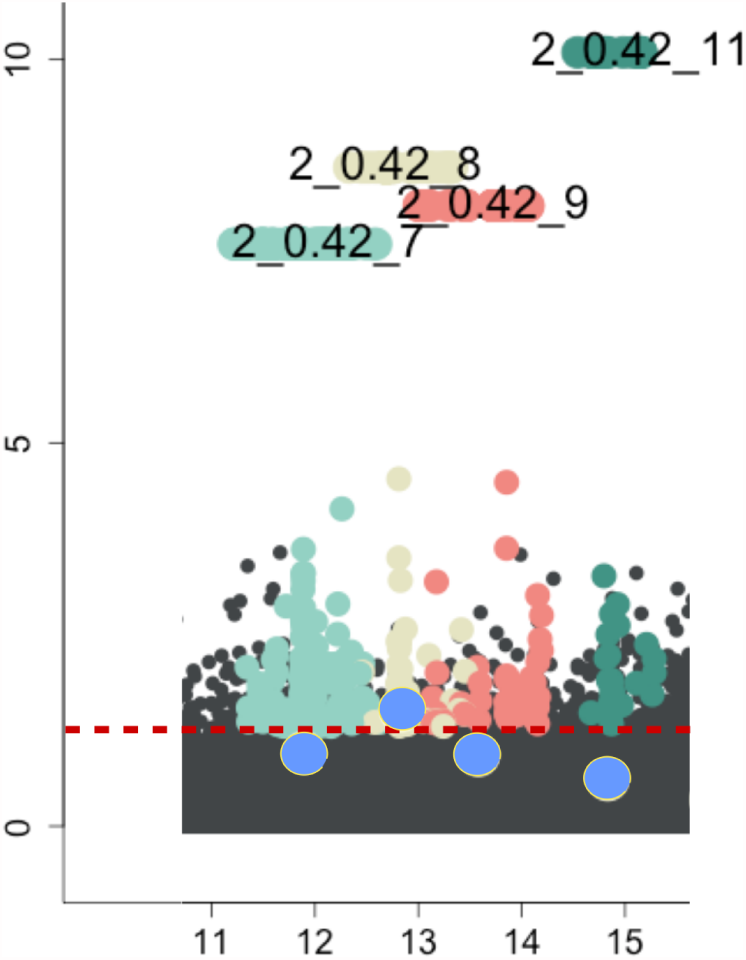
Example of putative false positive clusters in F20. The causative SNPs are marked in blue. Only region 2_0.42_8 contains the target of selection, but all other clusters contain hitchhiker SNPs only. The selected target is below the significance threshold (dashed red line) and therefore not included in the clustering.

### Experimental Data

The clustering of Barghi *et al.* (2019) is based on clustering parameters that were cross-checked with experimentally generated haplotypes. Hence, we were interested whether it is possible to recover the same clustering with haplovalidate, without consulting evolved haplotypes. Using the candidate SNPs from Barghi *et al.* (2019) with haplovalidate (MNCS 0.01) resulted in the identification of similar clusters. Instead of 88 clusters, haplovalidate detected 104 haplotype blocks of which all 104 overlap at least for 50 % with regions from Barghi *et al.* (2019). Vice versa, 70 regions detected by Barghi *et al.* (2019) overlap at least for 50% with haplovalidate (see figure 7).

**Figure 7:**
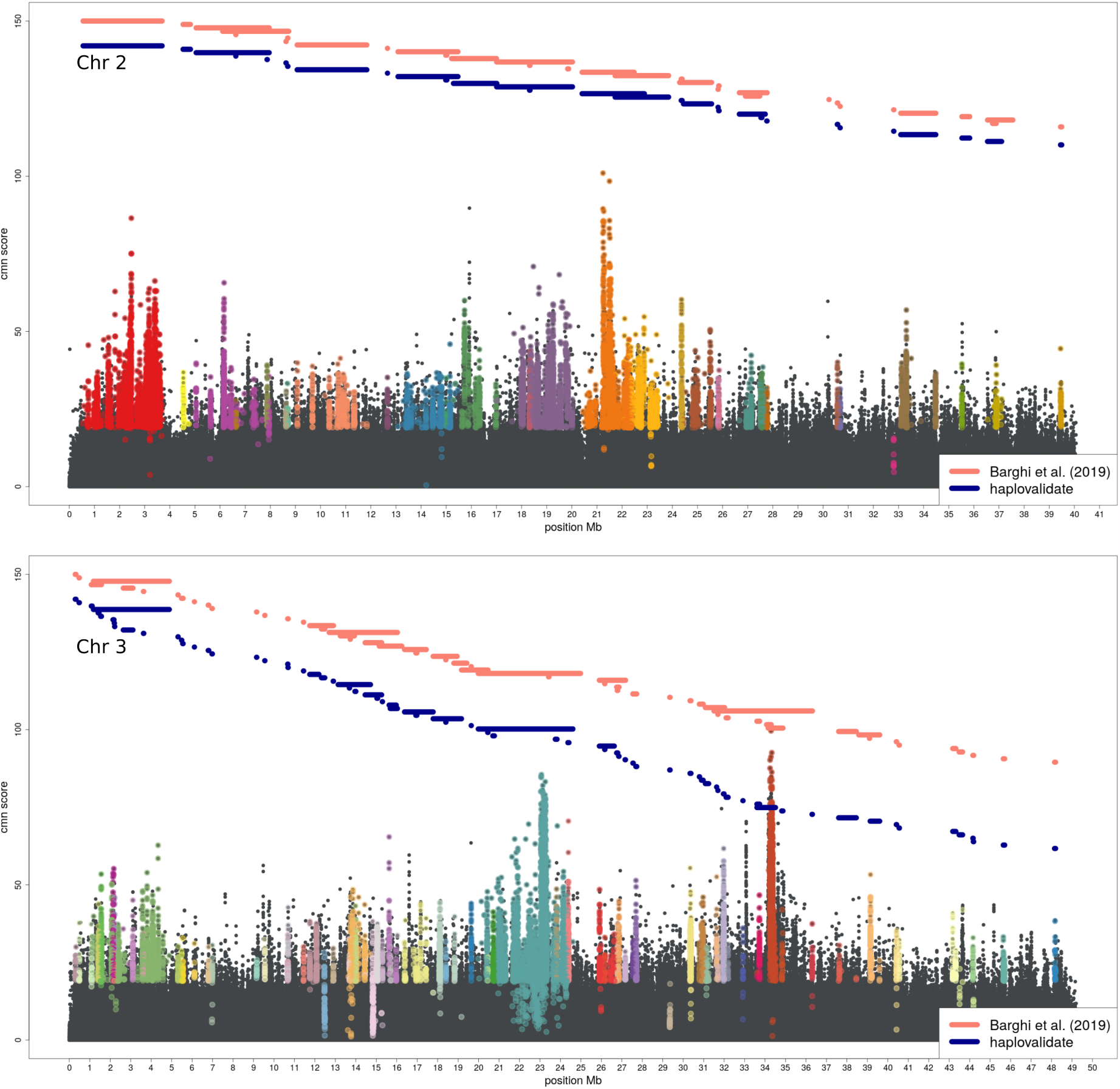
Manhattan plot of CMH p-values of Barghi *et al.* (2019) with SNPs belonging to clusters identified with haplovalidate shown in different colours. Pink lines indicate the genomic region spanned by haplotype blocks of Barghi *et al.* (2019) and blue lines represent those detected by haplovalidate.

When applied to the 1000 most significant SNPs for each chromosome of the data of Mallard *et al.* (2018), haplovalidate (MNCS 0.01) identified all five regions detected in the original study, including distinct regions containing the two top candidate genes (SNPs above dashed line in figure 8). Although both regions contain several genes, the two candidate genes SNF4 and Sestrin are among the genes with the highest number of outlier SNPs.

**Figure 8:**
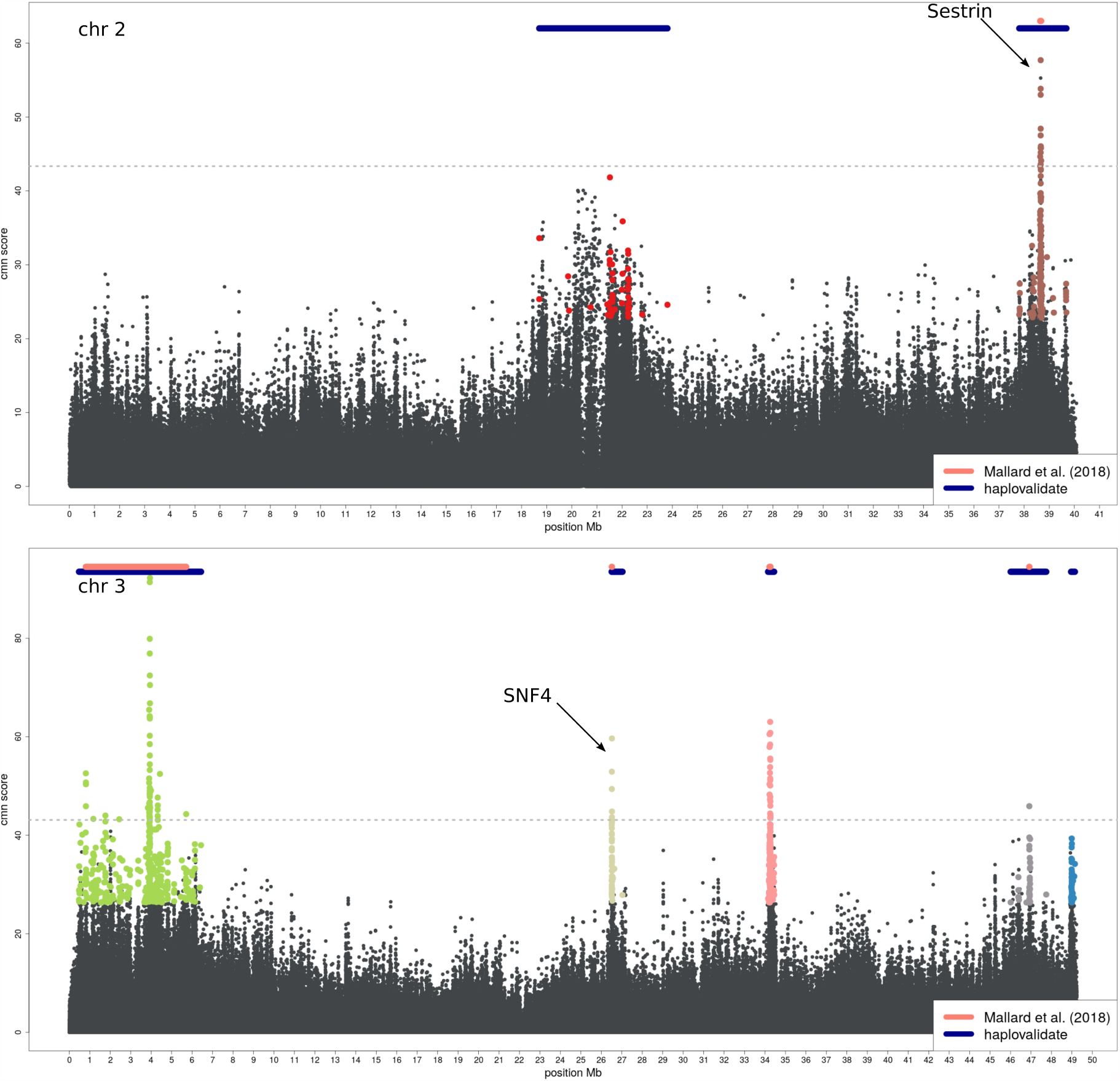
Manhattanplot showing the results of haplovalidate using the data of Mallard *et al.* (2018) on chromosome 2 (top) and chromosome 3 (bottom). Dots of the same colour represent SNPs belonging to the same reconstructed haplotype. The dashed line represents the threshold used in the original study. The two top candidate genes (Sestrin and SNF4) identified by Mallard *et al.* (2018) are top genes in two distinct regions.

## Discussion

Moving from a SNP-centric analysis to the identification of selected haplotype blocks provides a significant advancement of E&R studies (Barghi and Schlötterer, 2019). Introducing haplovalidate, which reconstructs selected haplotype blocks from genomic E&R time-series data, we provide a tool, to make the reconstruction of selected haplotypes a routine method that does not rely on the availability of haplotype information, neither from the founder population, nor from evolved individuals. Haplovalidate can be applied to a broad range of data, including few selection targets (Mallard *et al.*, 2018) as well as many (Barghi *et al.*, 2019). We attribute this generality to the data-driven selection of two key parameters of the clustering procedure, the minimum correlation between SNPs constituting a cluster and the window size.

Nevertheless, our study also demonstrated the limits of a haplotype block-based analysis of the adaptive architecture. We noted that a high fraction of the reconstructed haplotype blocks contained multiple selection targets. Interestingly, a similar observation was made by Sachdeva and Barton (2018) when analysing linked polygenic selection. In concordance with our study, the authors found that multiple-targets haplotypes out-competed other haplotypes over time.

The occurrence of multiple-target haplotypes leads to a substantial under-estimation of selection targets if each haplotype block is considered the outcome of selection operating on a single target in this block. Restricting the analysis to intermediate generations (up to F20 or F30) improved the resolution – significantly more single-target blocks were identified while fewer multiple-target blocks with a fewer selection targets are detected (see figure 5). Interestingly, most multiple-target haplotype blocks were already present in the founder population, indicating that the initial haplotype structure is an important factor for shaping the genomic signatures of selection. More work is needed to understand how the haplotype composition of the founder population in combination with the number of founder haplotypes affects the power of E&R studies.

## Material and Methods

We evaluated the performance of haplovalidate using a range of sweep simulations with different numbers of targets and selection coefficients. We applied haplovalidate to two already published *D. simulans* E&R studies and compared our results to the original studies. If not stated otherwise, all analyses were conducted with R version 3.6.0 (R Core Team, 2016) and the R package poolSeq version 0.3.1 (Taus *et al.*, 2017) and haploReconstruct 0.1.3_3 (Franssen *et al.*, 2017).

### Simulations

We performed 1000 genome-scale forward simulations covering the two main autosomes of *D. simulans* using using MimicrEE2 v206 (Vlachos and Kofler, 2018), simulating diploid sexual organisms and loci with constant selection coefficients. We mimicked an E&R experiment in *Drosophila simulans* consisting of 10 replicates with a population size of 1200 each, evolving for 60 generations. We extracted allele frequencies for every 10th generation (sync file format, Kofler *et al.* (2011)) and haplotypes for the most evolved generation (F60). We restricted our analysis to the main autosomes of *D. simulans* (chromosome 2 and chromosome 3).

The founder population was created from 189 experimentally phased haplotypes originating from a natural *D. simulans* population (Barghi *et al.*, 2019). We generated the same number of simulations for 16, 32, 64 or 128 selected loci (equal number of loci on both chromosomes). Starting allele frequencies and selection coefficients were taken from Barghi *et al.* (2019). The corresponding number of loci was drawn from the set of 99 selection targets without replacement. In the case of 128 loci, 29 randomly chosen value pairs were used twice. SNPs matching the allele frequency were randomly chosen from the founder population. We used the *D. simulans* specific recombination map (`Dsim_recombination_map_LOESS_100kb_1.txt`, Howie *et al.* (2019)). We generated “Pool-seq data” with 50x coverage and added sequencing noise by binomial sampling based on the allele frequencies.

### Candidate SNPs

Because only selected haplotypes are of interest, clustering is performed only on candidate SNPs, which change more in frequency than expected under neutrality. Candidate SNPs were identified by *χ*^2^-test and Cochran-Mantel-Haenszel (CMH) test accounting for drift and pool sequencing variance as implemented in the R package ACER (Spitzer *et al.*, 2019). All available time-points were used for calculating the test statistics and effective population size was calculated (estimateWndNe function from poolSeq package (Taus *et al.*, 2017)) for intermediate and most evolved generations. In addition, we corrected the p-values for multiple testing using the Benjamini-Hochberg method as implemented in the R function p.adjust. We chose candidates SNPs having a corrected p-value *<* 0.05 in the cmh-test or p-value *<* 0.001 in the *χ*^2^-test of any replicate. Using both tests allows for replicate specific responses, which could be missed by the cmh-test alone as it detects consistent changes across replicates. A more stringent p-value threshold for the *χ*^2^-test was used to include only replicate-specific candidates with a pronounced allele frequency change. If a simulation run had more than 50,000 candidate SNPs per chromosome, we randomly sampled 50,000 SNPs from the candidates to increase computational efficiency.

### Fine mapping of multiple-target haplotypes

We identified multiple-target haplotypes by comparing haplotype blocks reconstructed based all generations (up to 60) to those obtained from reconstructions using time points up to generation 20 (or generation 30 if not enough candidate SNPs were identified at F20). Regions with several haplotype blocks at the intermediate time point and only a single haplotype block in the later generation indicate a multiple-target haplotype.

The results of haplovalidate with and without intermediate generations were compared based on a each of the normalised summary statistics (i.e. true positive rate, single target fraction, multiple target fraction, multiple target per haplotype fraction, false positive rate, figure 5) using arcsine square-root transformation and the Welch’s t-test.

### Application to experimental data

We applied haplovalidate to two different *Drosophila* E&R data-sets which capture experimental evolution to a new temperature regime in *Drosophila simulans* but show different selection responses. Barghi *et al.* (2019) found 88 selected regions on the autosomes whereas Mallard *et al.* (2018) focused on two regions while analysing the top 100 cmh-test outlier SNPs. For an unbiased comparison of the clustering results, we aimed to use the same candidate SNPs as in the original studies. As the top 100 cmh-test outlier SNPs did not result in enough SNPs per window to perform haplotype reconstruction for the data of Mallard *et al.* (2018) we extended the SNP-set to the top 1000 outlier SNPs per chromosome.

### Availability

The R package haplovalidate is available at https://github.com/popgenvienna/haplovalidate. Summary statistics and scripts are available from the Dryad Digital Repository under XXX.

## Acknowledgements

Special thanks to Susanne Franssen for her help with haploReconstruct and her feedback. The authors also thank the members of the Institute of Population Genetics for discussion and support on the project and especially Anna Maria Langmüller and Robert Kofler for helpful comments on earlier versions of the manuscript. KAO was supported by a DFG Research Fellowship (OT 532/1-1). CS was supported by the European Research Council grant “ArchAdapt” and the Austrian Science Funds (FWF, P27630, P29133).

## References

Barghi, N. and Schlötterer, C. 2019. Shifting the paradigm in Evolve and Resequence studies: From analysis of single nucleotide polymorphisms to selected haplotype blocks. Molecular ecology, 28(3): 521–524.

Barghi, N., Tobler, R., Nolte, V., Jakšić, A. M., Mallard, F., Otte, K. A., Dolezal, M., Taus, T., Kofler, R., and Schlötterer, C. 2019. Genetic redundancy fuels polygenic adaptation in Drosophila. PLoS biology, 17(2): e3000128.

Burke, M. K., Dunham, J. P., Shahrestani, P., Thornton, K. R., Rose, M. R., and Long, A. D. 2010. Genome-wide analysis of a long-term evolution experiment with Drosophila. Nature, 467(7315): 587–590.

Burke, M. K., Liti, G., and Long, A. D. 2014. Standing genetic variation drives repeatable experimental evolution in outcrossing populations of saccharomyces cerevisiae. Molecular Biology and Evolution, 31(12): 3228–3239.

Cao, C. C. and Sun, X. 2015. Accurate estimation of haplotype frequency from pooled sequencing data and cost-effective identification of rare haplotype carriers by overlapping pool sequencing. Bioinformatics, 31(4): 515–522.

Cubillos, F. A., Parts, L., Salinas, F., Bergström, A., Scovacricchi, E., Zia, A., Illingworth, C. J., Mustonen, V., Ibstedt, S., Warringer, J., Louis, E. J., Durbin, R., and Liti, G. 2013. High-resolution mapping of complex traits with a four-parent advanced intercross yeast population. Genetics, 195(3): 1141–1155.

Fisher, R. A. 1915. Frequency Distribution of the Values of the Correlation Coefficient in Samples from an Indefinitely Large Population. Biometrika, 10(4): 507.

Franssen, S. U., Nolte, V., Tobler, R., and Schlötterer, C. 2015. Patterns of Linkage Disequilibrium and Long Range Hitchhiking in Evolving Experimental Drosophila melanogaster Populations. Molecular Biology and Evolution, 32(2): 495–509.

Franssen, S. U., Barton, N. H., and Schlotterer, C. 2017. Reconstruction of haplotype-blocks selected during experimental evolution. Molecular Biology and Evolution, 34(1): 174–184.

Griffin, P. C., Hangartner, S. B., Fournier-Level, A., and Hoffmann, A. A. 2017. Genomic trajectories to desiccation resistance: Convergence and divergence among replicate selected Drosophila lines. Genetics, 205(2): 871–890.

Howie, J. M., Mazzucco, R., Taus, T., Nolte, V., and Schlötterer, C. 2019. DNA Motifs Are Not General Predictors of Recombination in Two Drosophila Sister Species. Genome biology and evolution, 11(4): 1345–1357.

Jha, A. R., Miles, C. M., Lippert, N. R., Brown, C. D., White, K. P., and Kreitman, M. 2015. Whole-Genome Resequencing of Experimental Populations Reveals Polygenic Basis of Egg-Size Variation in Drosophila melanogaster. Molecular Biology and Evolution, 32(10): 2616–2632.

Kapun, M., Van Schalkwyk, H., McAllister, B., Flatt, T., and Schlötterer, C. 2014. Inference of chromosomal inversion dynamics from Pool-Seq data in natural and laboratory populations of Drosophila melanogaster. Molecular Ecology, 23(7): 1813–1827.

Kessner, D., Turner, T. L., and Novembre, J. 2013. Maximum likelihood estimation of frequencies of known haplotypes from pooled sequence data. Molecular Biology and Evolution, 30(5): 1145–1158.

Kofler, R. and Schlötterer, C. 2014. A Guide for the Design of Evolve and Resequencing Studies. Molecular Biology and Evolution, 31(2): 474–483.

Kofler, R., Pandey, R. V., and Schlötterer, C. 2011. PoPoolation2: Identifying differentiation between populations using sequencing of pooled DNA samples (Pool-Seq). Bioinformatics, 27(24): 3435–3436.

Langley, C. H., Crepeau, M., Cardeno, C., Corbett-Detig, R., and Stevens, K. 2011. Circumventing heterozygosity: Sequencing the amplified genome of a single haploid Drosophila melanogaster embryo. Genetics, 188(2): 239–246.

Long, A., Liti, G., Luptak, A., and Tenaillon, O. 2015. Elucidating the molecular architecture of adaptation via evolve and resequence experiments. Nature Reviews Genetics, 16: 567–582.

Long, Q., Jeffares, D. C., Zhang, Q., Ye, K., Nizhynska, V., Ning, Z., Tyler-Smith, C., and Nordborg, M. 2011. PoolHap: Inferring haplotype frequencies from pooled samples by next generation sequencing. PLoS ONE, 6(1): 1–7.

Mallard, F., Nolte, V., Tobler, R., Kapun, M., and Schlötterer, C. 2018. A simple genetic basis of adaptation to a novel thermal environment results in complex metabolic rewiring in Drosophila. Genome Biology, 19(1): 1–52.

Nuzhdin, S. V. and Turner, T. L. 2013. Promises and limitations of hitchhiking mapping. Current Opinion in Genetics & Development, 23(6): 694–699.

Orozco-terWengel, P., Kapun, M., Nolte, V., Kofler, R., Flatt, T., and Schlötterer, C. 2012. Adaptation of Drosophila to a novel laboratory environment reveals temporally heterogeneous trajectories of selected alleles. Molecular Ecology, 21(20): 4931–4941.

R Core Team 2016. R: A Language and Environment for Statistical Computing.

Remolina, S., Chang, P., and Leips, J. 2012. Genomic Basis of Aging and Life History Evolution in Drosophila melanogaster. Evolution, 66(11): 3390–403.

Sachdeva, H. and Barton, N. H. 2018. Replicability of introgression under linked, polygenic selection. Genetics, 210(4): 1411–1427.

Schlötterer, C., Tobler, R., Kofler, R., and Nolte, V. 2014. Sequencing pools of individuals — mining genome-wide polymorphism data without big funding. Nature Reviews Genetics, 15(11): 749–763.

Spitzer, K., Pelizzola, M., and Futschik, A. 2019. Modifying the Chi-square and the CMH test for population genetic inference: adapting to over-dispersion. BioArxiv.

Taus, T., Futschik, A., and Schlötterer, C. 2017. Quantifying Selection with Pool-Seq Time Series Data. Molecular biology and evolution, 34(11): 3023–3034.

Tenaillon, O., Rodríguez-Verdugo, A., Gaut, R. L., McDonald, P., Bennett, A. F., Long, A. D., and Gaut, B. S. 2012. The molecular diversity of adaptive convergence. Science, 335(6067): 457–461.

Tobler, R., Franssen, S. U., Kofler, R., Orozco-terWengel, P., Nolte, V., Hermisson, J., and Schlötterer, C. 2014. Massive Habitat-Specific Genomic Response in D. melanogaster Populations during Experimental Evolution in Hot and Cold Environments. Molecular Biology and Evolution, 31(2): 364–375.

Turner, T. L., Stewart, A. D., Fields, A. T., Rice, W. R., and Tarone, A. M. 2011. Population-Based Resequencing of Experimentally Evolved Populations Reveals the Genetic Basis of Body Size Variation in Drosophila melanogaster. PLoS Genetics, 7(3): e1001336.

Turner, T. L., Miller, P. M., and Cochrane, V. A. 2013. Combining genome-wide methods to investigate the genetic complexity of courtship song variation in Drosophila melanogaster. Molecular Biology and Evolution, 30(9): 2113–20.

Vlachos, C. and Kofler, R. 2018. MimicrEE2: Genome-wide forward simulations of Evolve and Resequencing studies. PLoS Computational Biology, 14(8): 1–10.

