## supplementary data for "A generalised approach to detect selected haplotype blocks in Evolve and Resequence experiments"

### S1 – Intermediate generation analysis for different numbers of selection targets

### S2 – False positive rate for early and late generations

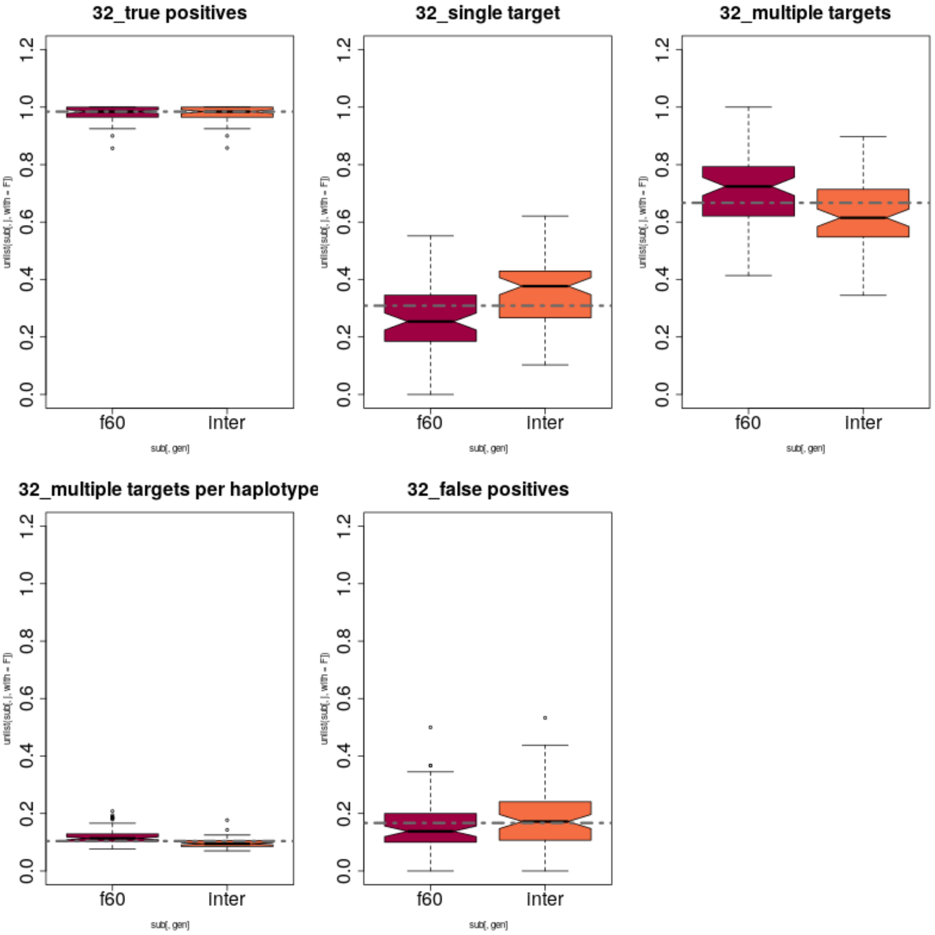


Figure 1: S1: Haplovalidate with and without including intermediate time-points for simulations containing 32 selection targets. A: True positive rate, B: fraction of selected regions with a single selection target, C: fraction of haplotype blocks with multiple targets, D average number of selection targets on a haplotype block, E: false positive rate. Dashed line represents the overall median for each parameter.

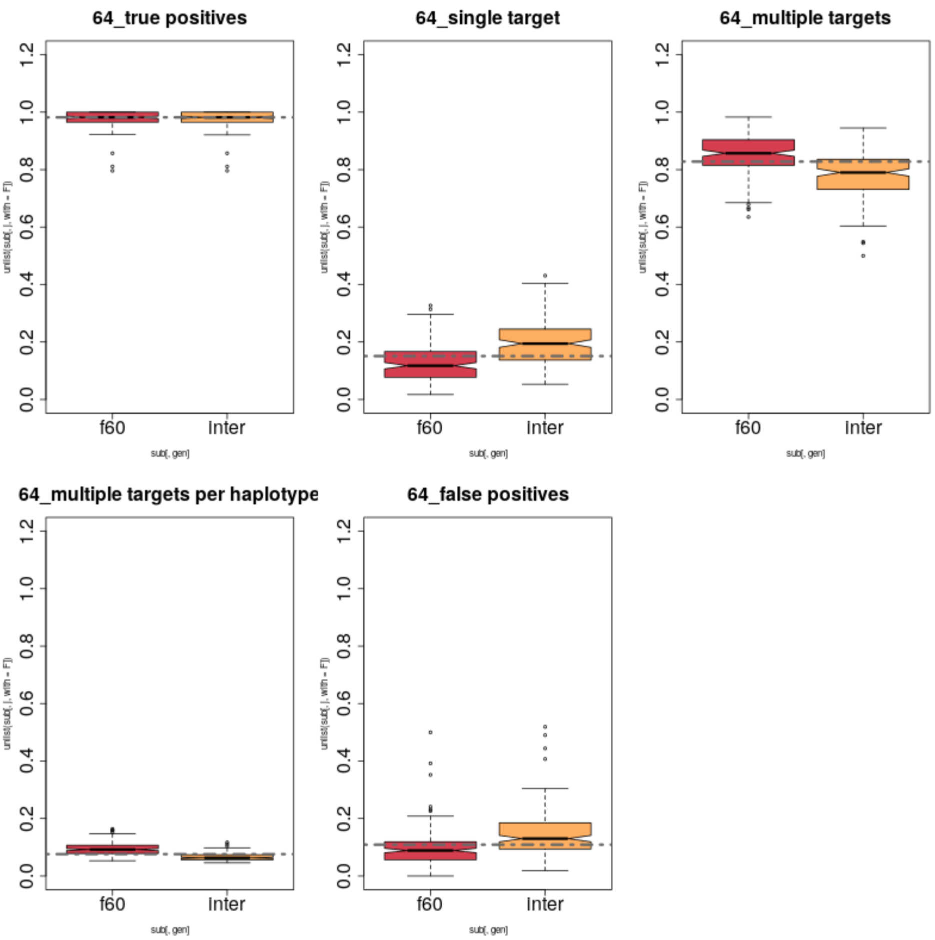


Figure 2: S1: Haplovalidate with and without including intermediate time-points for simulations containing 64 selection targets. A: True positive rate, B: fraction of selected regions with a single selection target, C: fraction of haplotype blocks with multiple targets, D average number of selection targets on a haplotype block, E: false positive rate. Dashed line represents the overall median for each parameter.

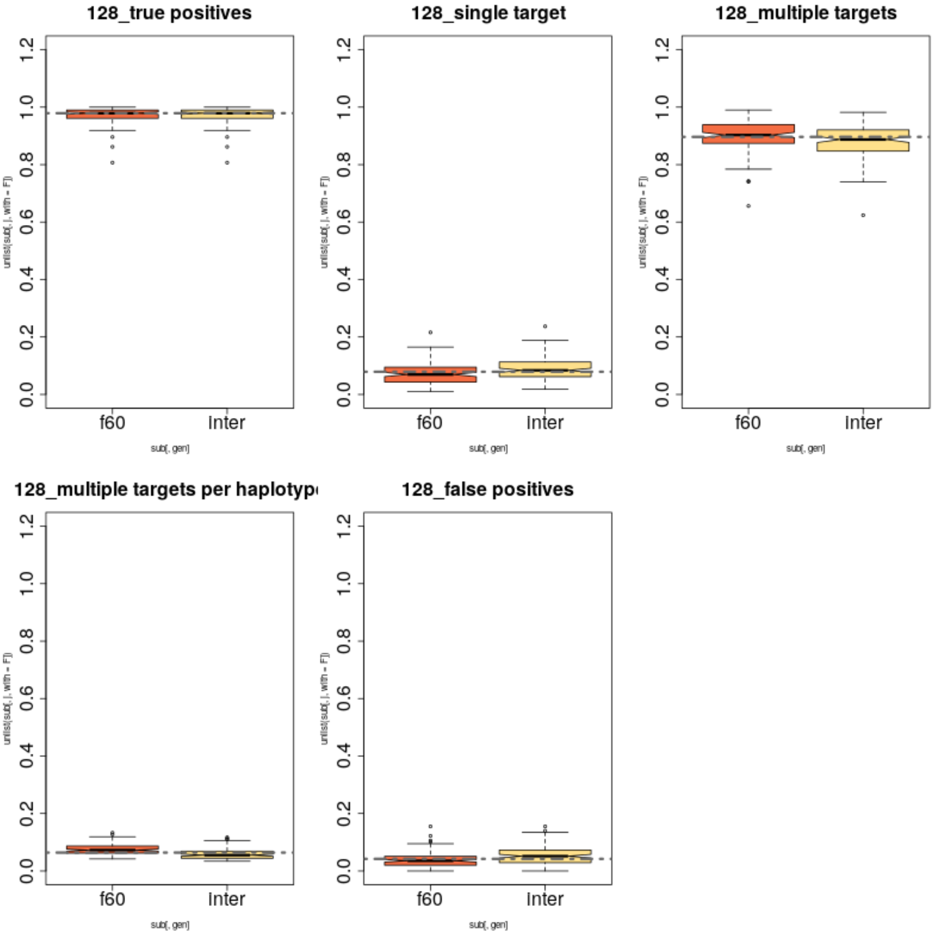


Figure 3: S1: Haplovalidate with and without including intermediate time-points for simulations containing 128 selection targets. A: True positive rate, B: fraction of selected regions with a single selection target, C: fraction of haplotype blocks with multiple targets, D average number of selection targets on a haplotype block, E: false positive rate. Dashed line represents the overall median for each parameter.

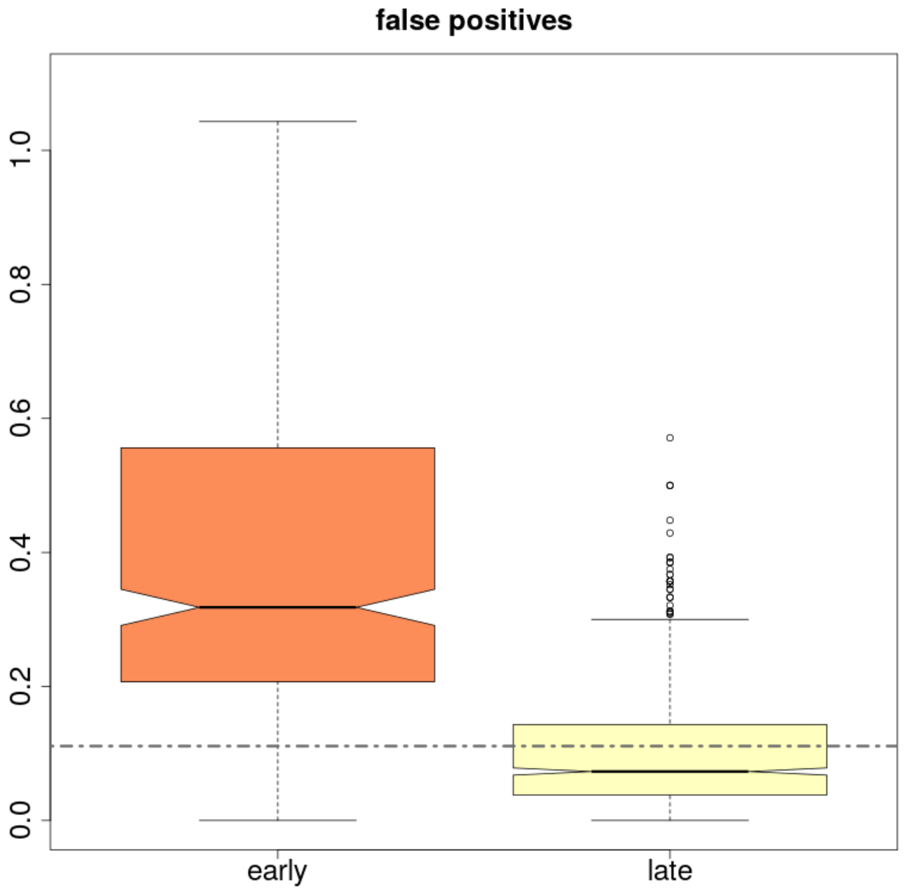


Figure 4: S2: False positive rates for the analysis of early (F20/F30) and late (F60) generations. Dashed line represents the overall median for each parameter.
